# “Temporal variability of the *Hyalomma marginatum*-borne pathogens in a sentinel site of the Occitanie region (France), a focus on the intriguing dynamics of *Rickettsia aeschlimannii*”

**DOI:** 10.1101/2024.04.16.589770

**Authors:** Joly-Kukla Charlotte, Stachurski Frédéric, Duhayon Maxime, Galon Clémence, Moutailler Sara, Pollet Thomas

## Abstract

Spatio-temporal scales have a clear influence on microbial community distribution and diversity and are thus essential to consider to identify and study the dynamics of microorganisms. The invasive tick species *Hyalomma marginatum* has recently become established in southern France. It carries pathogens of medical and veterinary interest including the Crimean-Congo hemorrhagic fever virus, *Rickettsia aeschlimannii, Theileria equi* and *Anaplasma phagocytophilum* among others. While the pathogenic communities of *H. marginatum* were identified and their spatial distribution characterised, their temporal dynamics are still unknown. We performed a monthly *H. marginatum* tick collection from February to September 2022 in a sentinel site in southern France in order to study their presence and temporal dynamics. On the 281 ticks analysed, we detected pathogens included *R. aeschlimannii, A. phagocytophilum* and *T. equi* with infection rates reaching 47.0%, 4.6% and 11.0% respectively. Overall, 14.6% of ticks were infected with at least *Theileria* or *Anaplasma*, with monthly fluctuations ranging from 2.9% to 28.6%. Strong temporal patterns were observed for each of the detected pathogens, particularly for *R. aeschlimannii* whose infection rates drastically increased at the beginning of summer, correlated with the monthly mean temperatures in the sentinel site. Based on these results, we hypothesized that *R. aeschlimannii* might be a secondary symbiont of *H. marginatum* that could play a role into stress response to temperature. The analysis of monthly and seasonal fluctuations in *H. marginatum*-borne pathogens allowed us to conclude that the risk of infection is present throughout *H. marginatum* activity period, but particularly in summer.

**Highlights:**

- Strong monthly fluctuations of pathogen infection rates were observed especially for *Rickettsia aeschlimannii*, currently identified as a human pathogen whose pathogenic status in humans and its symbiotic status in *H. marginatum* are both questioned.
- The increase in temperature is correlated with the increase in *Rickettsia aeschlimannii* infection rates, providing clues about its potential function as a symbiont in *H. marginatum*.
- Ticks are recurrently infected with at least one other pathogen belonging to *Theileria* or *Anaplasma* genera with monthly fluctuations ranging from 2.9% to 28.6%.
- The monthly dynamics of *H. marginatum*-borne pathogens are important to consider to assess the risk posed by this tick.

**Graphical abstract:** 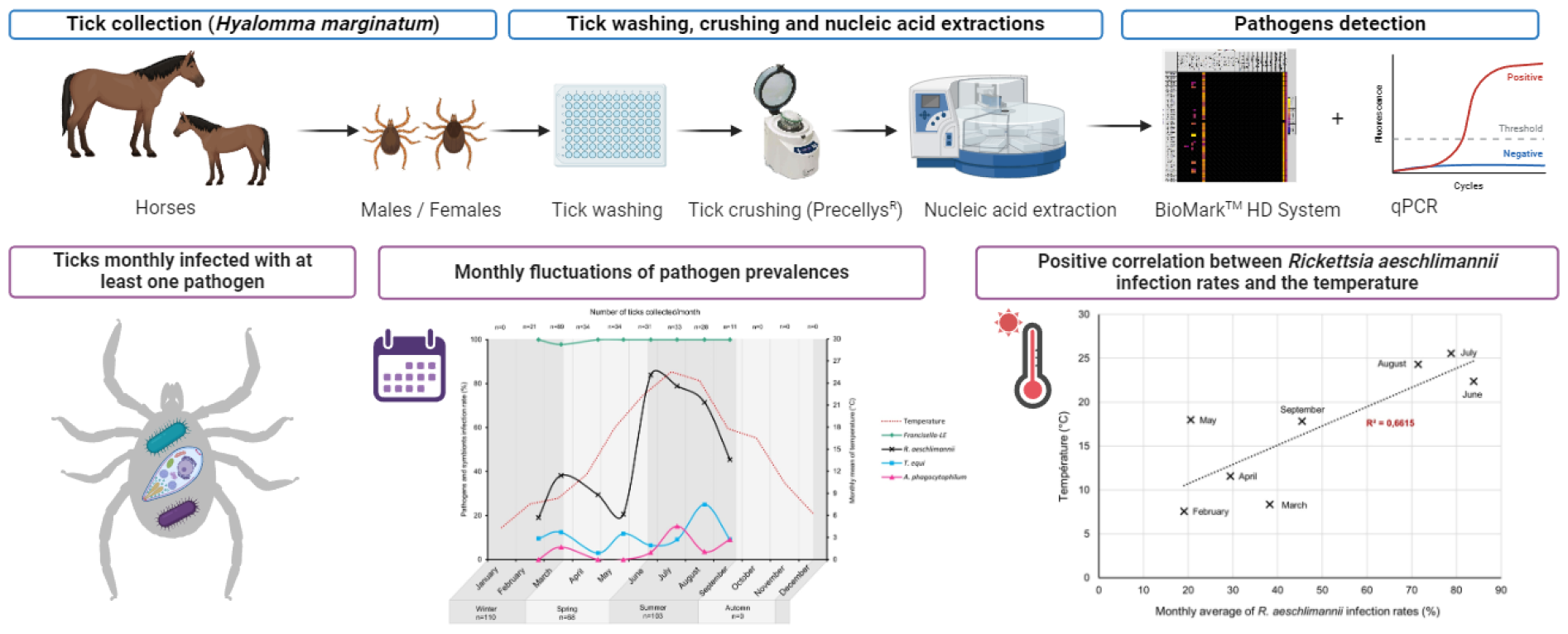

## 1 Introduction

The scale at which an epidemiological study is conducted has an undeniable effect on the interpretation of results. In the context of tick-borne diseases of medical and veterinary interest, the consideration of the spatial (tick location, tick organs) and the temporal scales (season, year) is essential to realise precise mappings of tick-borne pathogen (TBP) occurrence and dynamics (Pollet et al. 2020; Lejal et al. 2021; Lejal al. 2019a; Lejal et al. 2019b; Crowder et al. 2014; Halos et al. 2010; Joly-Kukla et al. 2024). These findings are crucial to consider for assessing the tick risk for both humans and animals.

Temporal patterns can influence both the tick life span and density due to environmental factors such as temperature, humidity, levels of canopy cover or forest fragmentation (Paul et al. 2016; Diuk-Wasser, VanAcker, and Fernandez 2021; Schulz, Mahling, and Pfister 2014; Randolph and Rogers 1997; Marchant et al. 2017; Wimberly et al. 2008). This variability influences the length of contact between ticks and their hosts, therefore impacting pathogen acquisition (Pollet et al. 2020). Longitudinal surveys are essential to visualise tick-borne pathogens occurrence fluctuations but also potential emergences, although they can be limited due to resource and time-consuming limitations (Sormunen et al. 2018). Several studies demonstrated how temporal trends affect the occurrence of pathogens from several tick species in different countries (Takken et al. 2017; Keesing et al. 2021; Tran et al. 2022; Foster et al. 2022; Kazimírová et al. 2023; Belkahia et al. 2017; Chvostáč et al. 2018; Coipan et al. 2013; Reye et al. 2010). In France, a three-years study on *Ixodes ricinus*-borne pathogens reported temporal fluctuations of prevalences of several common and less common TBP at the month, season and year scales (Lejal et al. 2019a). This study highlighted the limits of sporadic tick samplings and the need for a regular monitoring to determine pathogen prevalences.

In southern France, a new invasive tick species *Hyalomma marginatum*, recently established in the Mediterranean area (Vial et al. 2016; Stachurski and Vial 2018), is known to carry several animal and human pathogens such as *Rickettsia aeschlimannii, Theileria equi, Anaplasma phagocytophilum, Anaplasma marginale, Ehrlichia minasensis* and West Nile virus (Joly-Kukla et al. 2024; Bernard et al. 2024a). On the other hand, *H. marginatum* is known to be the reservoir and vector of the Crimean-Congo hemorrhagic fever virus, severely affecting humans (Bernard et al. 2022; Fillâtre, Revest, and Tattevin 2019; Hawman and Feldmann 2023). The virus was detected in southern France for the first time in ticks collected on cattle and horses in 2022 and 2023 (Bernard et al. 2024b).

The risk assessment linked to *H. marginatum* requires the knowledge of circulating pathogens and their infection rates but also the characterisation of their dynamics in both space and time. While an influence of the spatial scale was recently demonstrated on *H. marginatum* pathogen distribution (Joly-Kukla et al. 2024), the influence of the temporal scale has not been studied yet. As *H. marginatum* is an invasive tick species, the monitoring of its spread and the pathogens it carries need to implement a temporal surveillance over several years. Furthermore, since pathogen dynamics is possibly influenced by many environmental factors that vary through seasons, it is also essential to monthly assess the tick-borne pathogen dynamics. Such a temporal scale might also highlight the potential role of the bacterium *Rickettsia aeschlimannii*, previously detected with a high prevalence in *H. marginatum* ticks (87.3%) (Joly-Kukla et al. 2024). Even though this bacterium is currently identified as a human pathogen responsible for spotted fever, it is currently unclear if the strain detected in *H. marginatum* ticks represent the causative agent for the disease, especially given the lack of human cases (Beati et al. 1997; Parola, Paddock, and Raoult 2005; Portillo et al. 2015). In addition, there is no clear evidence about the real vectorial competency of *H. marginatum* for *R. aeschlimannii*. Finally, several recent studies hypothesize this bacterium might be a *H. marginatum* symbiont, whose role remains unknown and hypothetic. The analysis of seasonal trends in the occurrence of *R. aeschlimannii* could provide valuable insights into its function.

In this context, the objective of this study was thus to characterize the temporal patterns of *H. marginatum*-borne pathogens in a sentinel site in the south of France (Occitanie region).

## 2. Methods

### 2.1 Tick collection

The tick collection was carried out at a sentinel site called Mas de la Lauze, in Pompignan, located in the Gard department in southern France (**Figure 1**). This studied site covers a 200-hectare area of scrubland where horses, on which ticks are collected, live in semi-liberty. The tick collection occurred monthly between February and September 2022, during the activity period of *H. marginatum* adults, resulting in a total of 10 collection dates with three collection dates in March. We did not organise tick collection in January 2022 and between October and December 2022, as *H. marginatum* is not active during these periods (behavioural diapause). Collected ticks were morphologically identified using a binocular loupe and separated by sex. *H. marginatum* ticks were kept frozen at -80°C until further use. The dates of collection and the number of *H. marginatum* that were analysed were as followed: February 24^th^ (n=21 ticks), March 11^th^ (n=31), March 17^th^ (n=24), March 31^th^ (n=34), April 28^th^ (n=34), May 25^th^ (n=34), June 23^th^ (n=31), July 21^th^ (n=33), August 19^th^ (n=28) and September 16^th^ (n=11), resulting in a total of 281 *H. marginatum*. We chose to analyse a maximum of 34 ticks per date of collection while all the collected ticks were analysed when this number was below 34. A sex ratio (50/50) was respected when possible (in total 153 males and 128 females were analysed). From the different dates of tick collection, we defined three seasons as follow: winter: February 24^th^, March 11^th^, March 17^th^, March 31^th^; spring: April 28^th^, May 25^th^; summer: June 23^th^, July 21^th^, August 19^th^ and September 16^th^.

**Figure 1:**
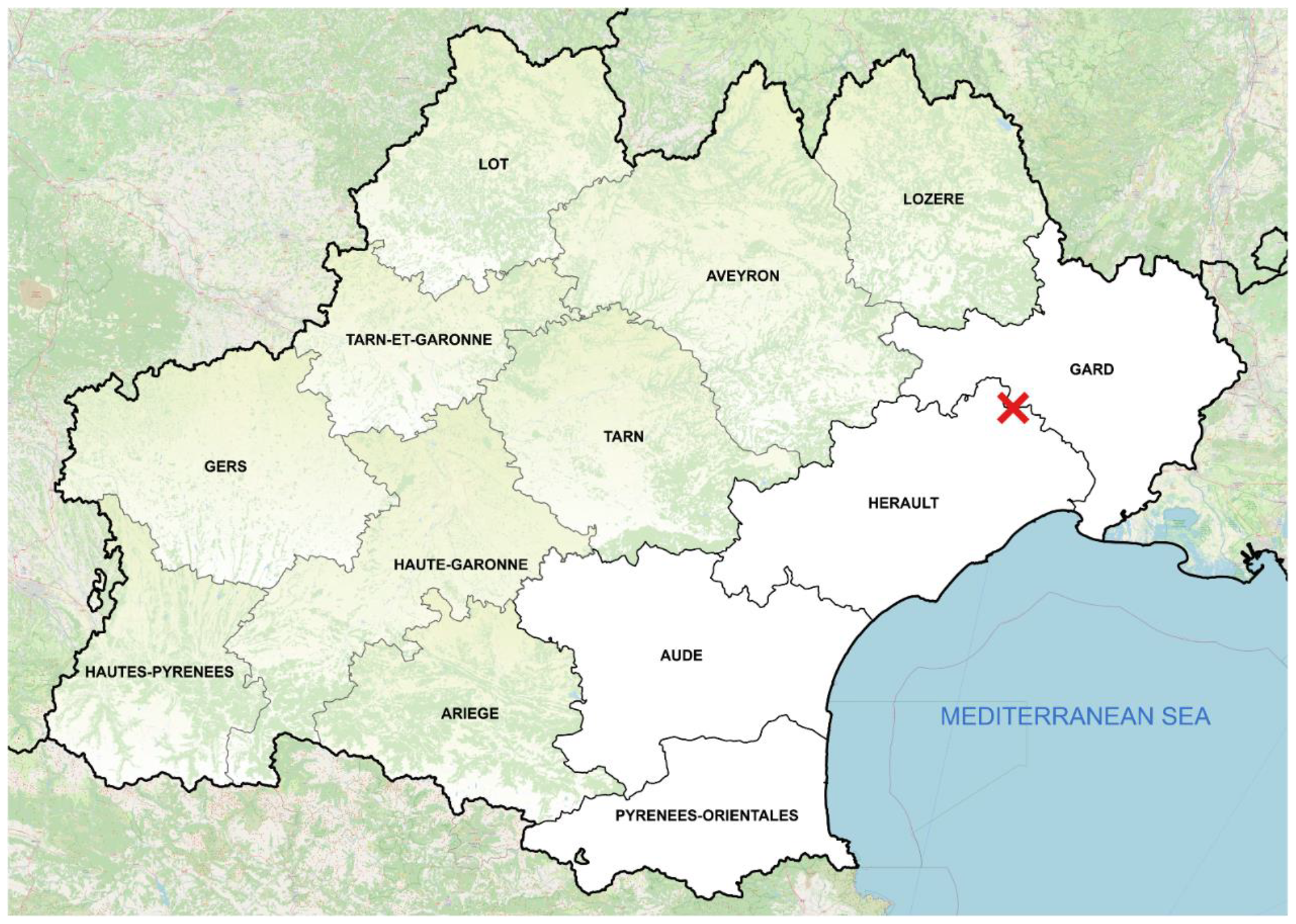
Map of the Occitanie region where the monthly tick sampling was conducted in the sentinel site “Pompignan” indicated by a red cross. Departments where *H. marginatum* populations are installed are coloured in white.

### 2.2 DNA and RNA extraction

Tick crushing and nucleic acids extraction were done in a BSL3 facility. Ticks were washed in bleach for 30 seconds then rinsed three times for one minute in milli Q water to eliminate the environmental microbes present on the tick cuticle (Binetruy et al. 2019). Ticks were then cut using a scalpel blade and crushed individually in the homogenizer Precellys®24 Dual (Bertin, France) at 5500 rpm for 40 sec, using three steel beads (2.8 mm, OZYME, France) in 400 µL of DMEM (Dulbecco’s Modified Eagle Medium, Eurobio Scientific France) with 10% foetal calf serum. Total DNA and RNA was extracted using the NucleoMag VET extraction kit (Macherey-Nagel, Germany) as described by the manufacturer’s instructions with the IDEAL™ 96 extraction robot (Innovative Diagnostics, France). Nucleic acids were eluted in 90 µL of elution buffer and stored at -20°C for DNA and -80°C for RNA until further analyses.

### 2.3 Tick-borne pathogens detection from tick DNA and RNA extractions

#### 2.3.1 Microfluidic PCR detection

##### 2.3.1.1 Reverse transcription

Samples were retrotranscribed in cDNA using 1 µL of extracted RNA with 1 µL of Reverse Transcription Master Mix and 3 µL of RNase-free ultrapure water provided with the kit (Standard Biotools, USA) using a thermal cycler (Eppendorf, Germany) with the following cycles: 5 min at 25°C, 30 min at 42°C and 5 min at 85°C with a final hold at 10°C.

##### 2.3.1.2 Targeted tick-borne pathogens

A design of 48 sets of primers/probes was used to target tick-borne pathogens (bacteria, parasites and viruses) commonly reported in *H. marginatum*. Please refer to Joly-Kukla et al. 2024 for the list of the targeted genes.

##### 2.3.1.3 Pre-amplification

Each sample was pre-amplified using 1.25 µL of DNA mix (1:1 volume ratio of DNA and cDNA) with 1 µL of the PreAmp Master mix (Standard Biotools, USA), 1.5 µL of ultra-pure water and 1.25 µL of the pooled designs. PCRs were performed using a thermal cycler with the following cycles: 2 min at 95°C, 14 cycles at 95°C for 15 sec and 60°C for 4 min and finally a 4-min hold at 4°C. A negative control was used for each plate with ultra-pure water. Amplicons were diluted 1:10 with ultra-pure water and stored at -20°C until further use.

##### 2.3.1.4 BioMark™ assay

The BioMark™ real-time PCR system (Standards Biotools, USA) was used for high-throughput microfluidic real-time PCR amplification using the 48.48 dynamic array (Standard Biotools, USA). The chips dispense 48 PCR mixes and 48 samples into individual wells, after which on-chip microfluidics assemble PCR reactions in individual chambers prior to thermal cycling resulting in 2,304 individual reactions. In one single experiment, 47 ticks and one negative control are being tested. For more details, please see (Gondard et al. 2018; 2020; Michelet et al. 2014; Joly-Kukla et al. 2024).

##### 2.3.1.5 Validation of the results by PCR and sequencing

Conventional PCRs or qPCR were then performed for TBP positive-samples using different sets of primers than those used in the BioMark™ assay to confirm the presence of pathogenic DNA, please refer to Joly-Kukla et al. 2024 for more details. Amplicons were sequenced by Azenta Life Sciences (Germany) using Sanger-EZ sequencing and assembled using the Geneious software (Biomatters, New-Zealand). An online BLAST (National Center for Biotechnology Information) was done to compare results with published sequences in GenBank sequence databases.

#### 2.3.2 Detection and quantification of *T. equi* and *R. aeschlimannii* by duplex Real-Time Fluorescence Quantitative PCR

Tick samples were also screened for the detection and quantification of *T. equi* and *R. aeschlimannii* using a qPCR approach with primers and probes targeting different genes than those used in the BioMark™ assay, i.e. the 18S rRNA gene and the *Ompb* gene respectively, please see Joly-Kukla et al. 2024 for more details. Five different genotypes of *T. equi* (designated A-E) are known to circulate in Europe. We thus wanted to be sure that we could detect all of those five genotypes if present (Nagore et al. 2004; Bhoora et al. 2009; Salim et al. 2010; Qablan et al. 2013). The Takyon™ No ROX Probe 2X MasterMix Blue dTTP (Eurogentec, Belgium) was used with a final reaction volume of 20 µL containing 10 µL of Master Mix 2X (final concentration 1X), 5 µL of RNase free water, 1 µL of each primer (0.5 µM), probes (0.25 µM), and 2 µL of DNA template. The reaction was carried out using a thermal cycler according to the following cycles: 3 min at 95°C, 45 cycles at 95°C for 10 sec, 55°C for 30 sec and 72°C for 30 sec. Positive controls for both *R. aeschlimannii* and *T. equi* were prepared using a recombinant plasmid from the TA cloning® kit (Invitrogen, USA). A 10-fold serial dilution of the plasmid (from an initial concentration of 0.5×10^8^ copy number/μL) was used to generate standard positive plasmids from 2.5×10^5^ copy number/μL to 2.5×10^−1^ copy number/μL. Samples were detected in duplicates and quantified using the standard plasmids. For *R. aeschlimannii*, we considered negative samples whose Cq number was higher than Cq 37. This detection limit was established regarding the last dilution of the standard curve that could be detected by qPCR. For *T. equi*, most samples were close to or below the detection limit established with the *T. equi* standard curve. Because the protozoan *T. equi* may be circulating at low levels, we included all positive samples.

### 2.4 Statistical analyses

The multivariate analysis was performed with R software 4.3.0. Generalized linear models (R package Lme4 (Bates et al. 2015)) were used to test the effect of both the season (winter/spring/summer), and the tick sex (male/female) on the infection rates of *R. aeschlimannii, T. equi, A. phagocytophilum* and *Francisella*-LE (presence/absence, binomial distribution). We decided to include *Francisella*-LE in the analysis as a control since this bacterium is identified as a *H. marginatum* primary endosymbiont (Duron et al. 2018; 2017; Azagi et al. 2017). A gamma distribution was used for *R. aeschlimannii* loads. Model assessment was based on Akaike’s information criterion (AIC). Significance of variables has been assessed using the ‘ANOVA’ procedure within the package ‘car’ which performs a type III hypothesis (Fox and Weisberg 2018). The post-hoc tests were conducted using the function ‘emmeans’ (Tukey HSD test). The data used for the analyses were generated using the BioMark™ data for *A. phagocytophilum* and using the qPCR data for *R. aeschlimannii* and *T. equi*.

The correlation between *R. aeschlimannii* and the temperature was performed using a linear regression. A Spearman correlation analysis was then performed using the R software to statistically analyse the significance of the correlation.

## 3. Results

### 3.1 Detected microorganisms and their global infection rates *in H. marginatum*

Among all analyzed ticks collected from February to September, 132 tick samples were detected positive for *R. aeschlimannii* by qPCR (*OmpB* gene) and 125 samples using the BioMark™ assay (*glta* gene). The presence of *R. aeschlimannii* was confirmed by the sequencing of the *OmpB* gene with 100% homology and one sequence was deposited in GenBank (AN: PP663276). The infection rates estimated by the qPCR assay (*OmpB* gene) and the BioMark™ assay (*glta* gene) were respectively 47.0% [41.1 – 52.8%] and 44.5% [38.6 – 50.3%]. The qPCR data were selected for the statistical analysis of *R. aeschlimannii*.

Concerning the other pathogens, 14.6% [10,4 – 18,7%] of the ticks were infected with at least *Theileria equi* or *Anaplasma phagocytophilum*. In details, 31 ticks were positively detected for *T. equi* by qPCR, resulting in an infection rate of 11.0% [7.3-14.7%]. Among pathogens belonging to the genus *Anaplasma*, eight ticks were positive for *Anaplasma phagocytophilum* alone and five for both *A. phagocytophilum* and *Anaplama* spp. designs. While a specific band corresponding to the *msp2* gene of *A. phagocytophilum* was visible in the gel, no readable sequences were obtained after sending amplicons from confirmation PCRs. However, please note that the 13 samples positive for either *A. phagocytophilum* alone or both designs were considered infected by this bacterium since *A. phagocytophilum* was detected in ticks the same year from the same geographic area and confirmed by classical PCR in another study (Joly-Kukla et al. 2024) with an infection rate reaching 4.6% in average [2.2 – 7.1%].

Finally, the endosymbiont *Francisella*-LE was detected in 279 samples using the BioMark™ assay and confirmed by a specific *Francisella*-LE qPCR (Duron et al. 2018). *Francisella*-LE infection rate was 99.3% in average [98.3 – 100%].

Among all collected ticks, 1.1% [0 – 2.3%] were co-infected with *A. phagocytophilum* and *T. equi. Francisella*-LE was not considered in co-infection since its status as a primary endosymbiont, neither *R. aeschlimannii* whose pathogenic status will be discussed. Please finally note that no ticks were infected by any of the viruses tested.

### 3.2 Temporal patterns in *H. marginatum*-borne pathogens

From monthly collected ticks (n=281), we determined temporal patterns of each *Hyalomma marginatum*-borne pathogens (**Figure 2, Table 1, Table 2)**. Ticks were recurrently infected with at least one pathogen (*T. equi* or *A. phagocytophilum*) every month, with a minimum of 2.9% of ticks infected in April to a maximum of 28.6% in August. At the seasonal scale, 2.6 times more ticks were infected with at least one pathogen in summer (19.4% [11.6 -27.2%]) compared to spring (7.4% [1.0 – 13.7%]). Interestingly, this percentage was quite high in winter: 14.5% [7.9 – 21.2%].

**Table 1:**
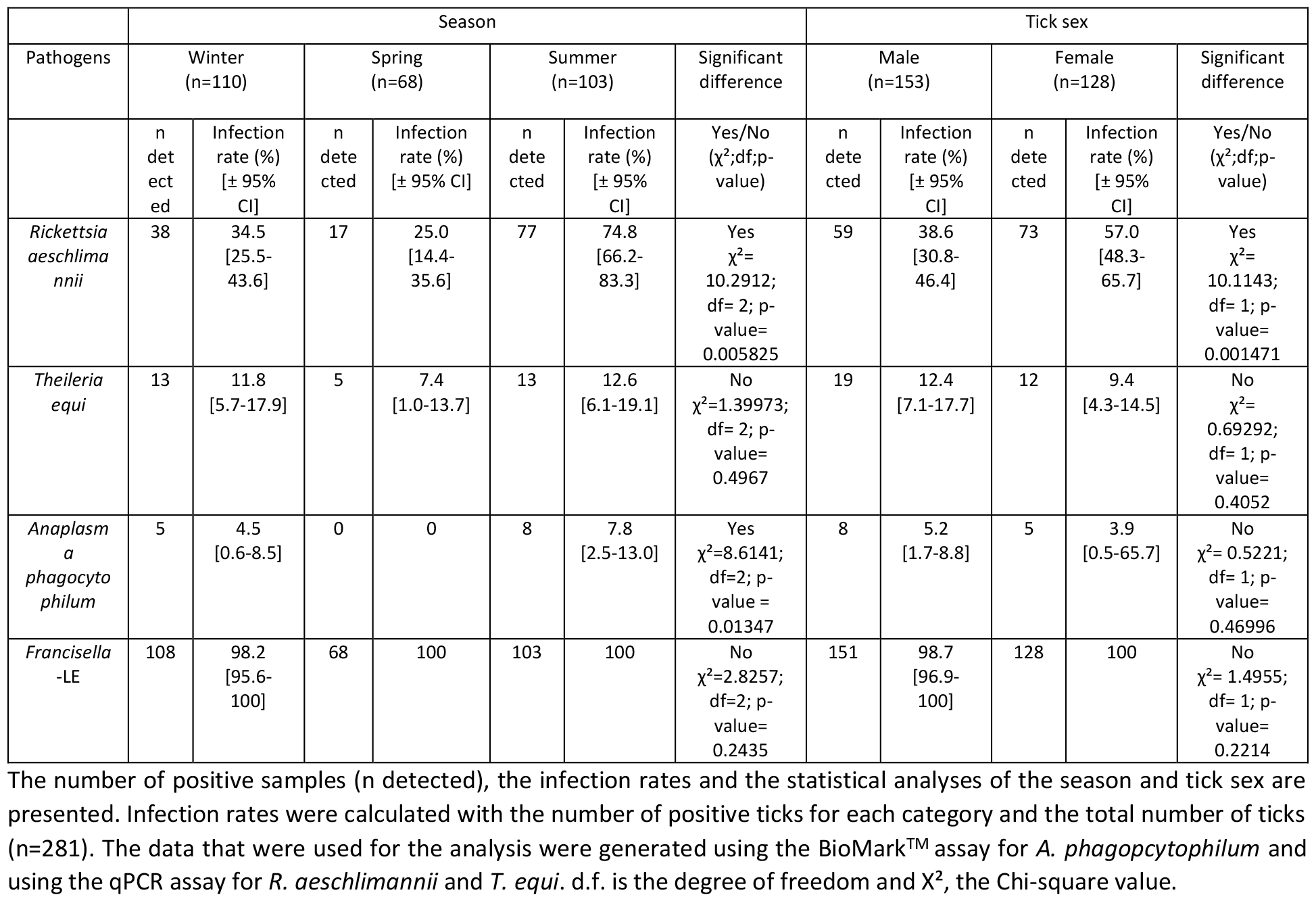
Multivariate analysis of *H. marginatum-*borne pathogens according to the season and the tick sex.

| Pathogens | Season |  |  |  |  |  |  | Tick sex |  |  |  |  |
| --- | --- | --- | --- | --- | --- | --- | --- | --- | --- | --- | --- | --- |
|  | Winter (n=110) |  | Spring (n=68) |  | Summer (n=103) |  | Significant difference | Male (n=153) |  | Female (n=128) |  | Significant difference |
| | n detected | Infection rate (%) [± 95% CI] | n detected | Infection rate (%) [± 95% CI] | n detected | Infection rate (%) [± 95% CI] | Yes/No ($\chi^2$;df;p-value) | n detected | Infection rate (%) [± 95% CI] | n detected | Infection rate (%) [± 95% CI] | Yes/No ($\chi^2$;df;p-value) |
| *Rickettsia aeschlimannii* | 38 | 34.5 [25.5-43.6] | 17 | 25.0 [14.4-35.6] | 77 | 74.8 [66.2-83.3] | Yes<br>$\chi^2=10.2912$; df= 2; p-value= 0.005825 | 59 | 38.6 [30.8-46.4] | 73 | 57.0 [48.3-65.7] | Yes<br>$\chi^2=10.1143$; df= 1; p-value= 0.001471 |
| *Theileria equi* | 13 | 11.8 [5.7-17.9] | 5 | 7.4 [1.0-13.7] | 13 | 12.6 [6.1-19.1] | No<br>$\chi^2=1.39973$; df= 2; p-value= 0.4967 | 19 | 12.4 [7.1-17.7] | 12 | 9.4 [4.3-14.5] | No<br>$\chi^2=0.69292$; df= 1; p-value= 0.4052 |
| *Anaplasma phagocytophilum* | 5 | 4.5 [0.6-8.5] | 0 | 0 | 8 | 7.8 [2.5-13.0] | Yes<br>$\chi^2=8.6141$; df=2; p-value= 0.01347 | 8 | 5.2 [1.7-8.8] | 5 | 3.9 [0.5-65.7] | No<br>$\chi^2=0.5221$; df= 1; p-value= 0.46996 |
| *Francisella*-LE | 108 | 98.2 [95.6-100] | 68 | 100 | 103 | 100 | No<br>$\chi^2=2.8257$; df=2; p-value= 0.2435 | 151 | 98.7 [96.9-100] | 128 | 100 | No<br>$\chi^2=1.4955$; df= 1; p-value= 0.2214 |

The number of positive samples (n detected), the infection rates and the statistical analyses of the season and tick sex are presented. Infection rates were calculated with the number of positive ticks for each category and the total number of ticks (n=281). The data that were used for the analysis were generated using the BioMark™ assay for *A. phagocytophilum* and using the qPCR assay for *R. aeschlimannii* and *T. equi*. d.f. is the degree of freedom and $\chi^2$ , the Chi-square value.

**Table 2:**
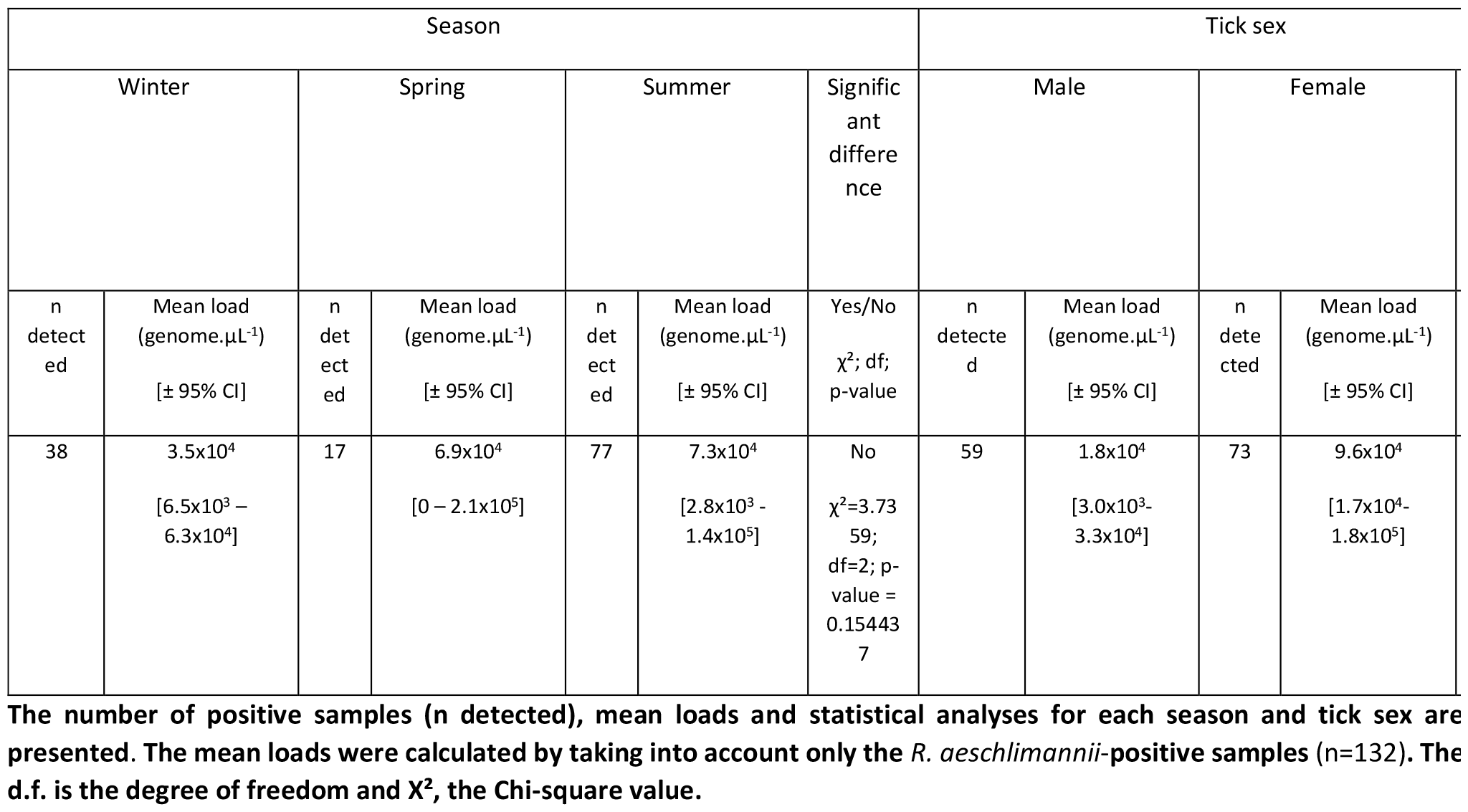
Multivariate analysis of *R. aeschlimannii* loads according to the season and the tick sex.

**Figure 2:**
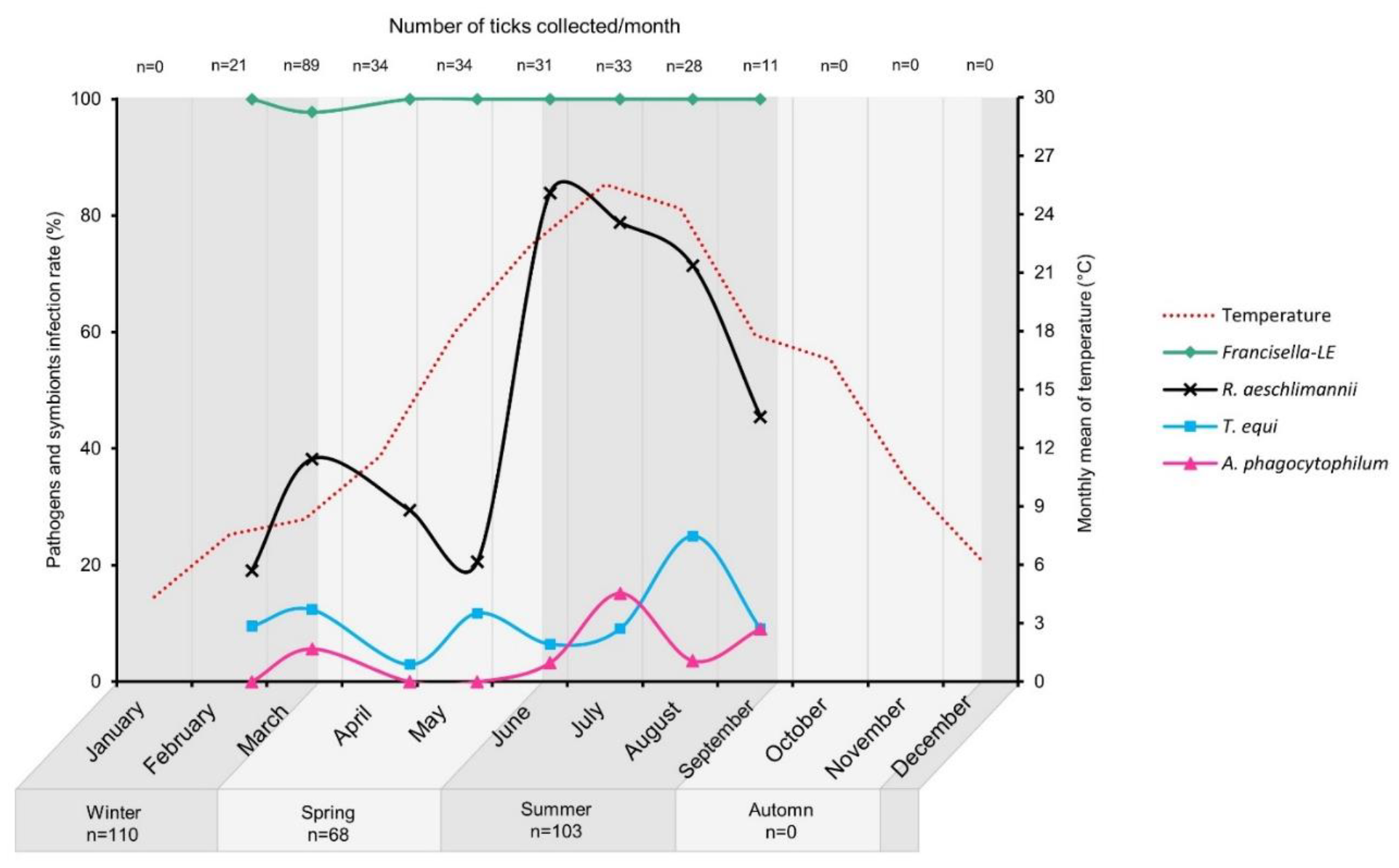
Infection rates (%) of *H. marginatum* pathogens and symbionts for each monthly date of collection (represented by shaped-dots). The number of collected ticks (abbreviation n) per month is indicated above the figure and below the figure for the seasons. Winter: February to March; Spring: April to May; Summer: June to September. The monthly mean temperature (°C) of the collection site Pompignan obtained through the R package Open-Meteo (https://open-meteo.com/) is plotted.

*R. aeschlimannii* infection rates were ranged from 19.0% in February to 83.9% in June. In addition, a drastic increase was reported between May: 20.6% [6.3 – 34.9%] and the following month June: 83.9% [70.2 – 97.6%]. Monthly infection rates in summer were all higher (74.8% in average [66.2 – 83.3%]) than those estimated in winter in average 34.5% [25.5 – 43.6%] and spring 25.0% in average [14.4 – 35.6%]. Indeed, the infection rates in summer were statistically higher compared to winter (χ^2^= 10.2912; df= 1; p-value= 0.005825) (**Table 1, Figure 2**). We also compared the variation of *R. aeschlimannii* infection rates that were assessed using two different targeted genes and techniques BioMark™ assay (target gene: citrate synthase gene), and the qPCR approach targeting the *OmpB* gene. The variation (from 0 to 3.2%) was limited for most of months (March, April, May, June and September). Bigger variations were observed for February, July and August (from 12.1 to 17.9%) although monthly trends were similar using the two techniques and targeted genes (**Figure S1**). As temporal patterns affecting *H. marginatum* pathogens can be directly linked to abiotic conditions, the monthly average of temperature (°C) in the sentinel site was plotted (**Figure 2**). The linear regression between the monthly infection rates of *R. aeschlimannii* and temperature revealed a significant positive correlation (Spearman ρ= 0.7619, p-value= 0.01838) (**Figure 3**).

**Figure 3:**
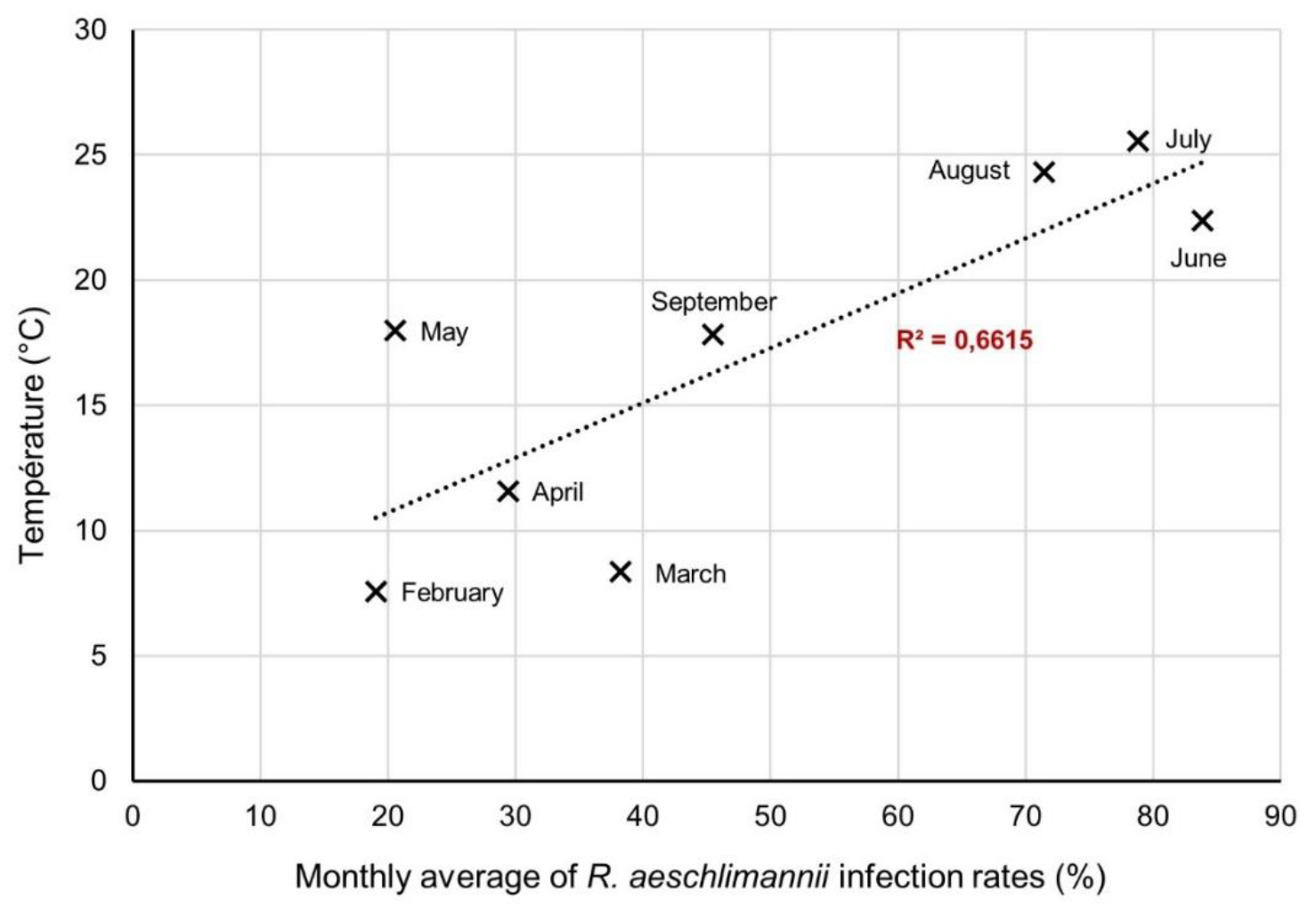
Linear correlation analysis of the monthly average of *R. aeschlimannii* infection rates (%) and the monthly temperature average (°C).

Beyond the seasonal influence, we also observed that infections rates were significantly different between males and females (χ^2^= 10.1143; df= 1; p-value= 0.001471) (**Table 1**). Indeed, infection rates during the whole period from February to September were significantly higher in females: 57.0% in average [48.3-65.7%] compared to males: 38.6% in average [30.8-46.4%]. Finally, the interaction between both the season and the tick sex variables was also significant (χ^2^= 9.5131; df= 2; p-value= 0.008595). Based on these results for *R. aeschlimannii*, we assessed the impact of the season in both males and females separately (**Figure 4A**). *R. aeschlimannii* infection rate in females was significantly higher in summer (71.7% [59.9-83.4%]) compared to spring (41.2% [23.7-58.6%]) (p-value= 0.0122). For information, its infection rate in winter was 47.1% [29.4-64.7%]. In male ticks, *R. aeschlimannii* was significantly higher in summer: 79.1% [66.4-91.7%] (p-value< 0.0001) compared to winter: 28.9% [18.5-39.4%] and spring: 8.8% [0-18.9%] (p-value< 0.0001). While the season did not have a significant influence on *R. aeschlimannii* loads (**Table 2**), the loads were significantly different between males and females (χ^2^= 9.7068; df= 1; p-value= 0.001836), with higher loads in females: 9.6x10^4^ [1.7x10^4^-1.8x10^5^] compared to males: 1.8x10^4^ [3.0x10^3^-3.3x10^4^] (p-value= 0.0018) (**Figure 4B, Table 2**).

**Figure 4:**
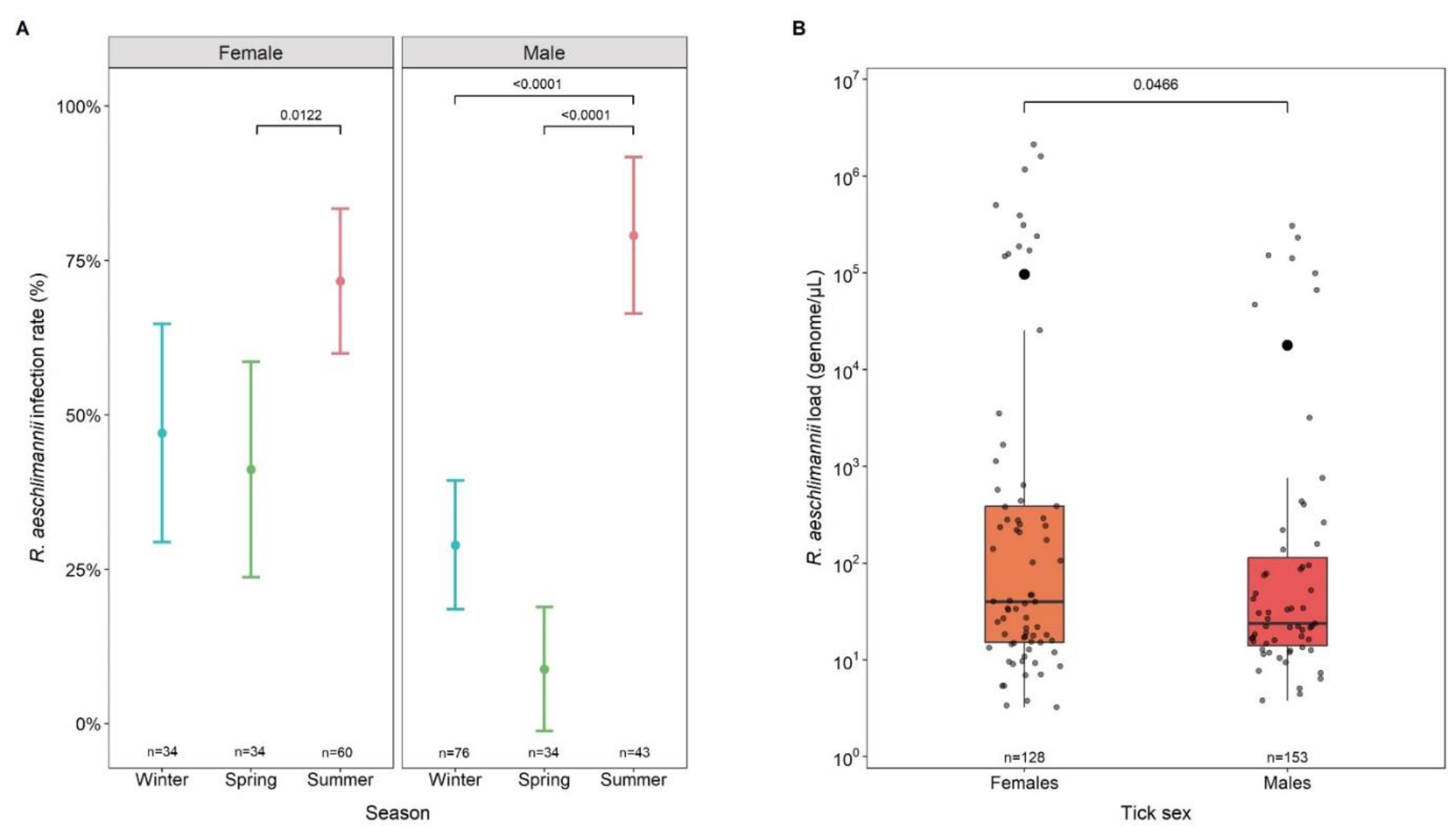
Seasonal trends of *R. aeschlimannii*. **A**. The average seasonal infection rate in % is represented by a dot with the confidence interval. **B**. The bacterial loads obtained by qPCR in genome.µL^-1^ are represented for each tick by grey dots and the mean is symbolized by the black dots. The boxplots summarise the median, 1st and 3rd quartiles. The number of collected ticks (abbreviation n) is indicated above each category. Winter: February to March; Spring: April to May; Summer: June to September. Only significant differences and the associated p-values (post-hoc test) are presented.

Interestingly, the difference between the lowest and the highest monthly infection rate for *R. aeschlimannii* was higher in males: 94.1% (from 5.9% in May, n=17 to 100% in June, n=14) compared to females: 52.9% (from 35.3% in May, n=17 to 88.2% in August, n=17) (**Figure S2A**). Finally, 0% infection rates were observed for females in February and males in September.

The presence of *T. equi, A. phagocytophilum* and *Francisella*-LE was not influenced by the tick sex (**Table 1, Figure S2**). The infection rates of *T. equi* were variable between months (from 2.9% in April to 25.0% in August), although no significant differences between the seasons were observed. However, for *A. phagocytophilum* a significant difference was reported according to the season (χ^2^= 8.6141; df= 2; p-value= 0.01347) (**Table 1**). Its infection rates were ranged from 0% in spring to 7.8% in average [2.5 – 13.0%] in summer and 4.5% in average [0.6 – 8.5%] in winter. Finally, the primary symbiont *Francisella*-LE was detected in almost all collected ticks (279/281) regardless of the season (**Table 1**).

## 4 Discussion

*Hyalomma marginatum* recently established in the south of France is known to potentially transmit several human and animal pathogens and thus represent a major concern for public and animal health. Assessing the risk represented by this tick species firstly requires to identify the *H. marginatum*-borne pathogen composition and characterize their temporal dynamics. In this context, we thus examined tick-borne pathogen (TBP) dynamics in *H. marginatum* collected monthly from February to September 2022 in a sentinel site in south of France.

### 4.1 *Hyalomma marginatum*-borne pathogen composition and prevalences

Three major pathogens have been detected in the 281 collected *Hyalomma marginatum*: *R. aeschlimannii* (47.0%), *T. equi* (11.0%) and *A. phagocytophilum* (4.6%). Overall, although the vector competence of *H. marginatum* for these pathogens is still unclear, they are already known to circulate in the Occitanie region (Joly-Kukla et al. 2024; Bernard et al. 2024a). 14.6% of ticks were infected with at least *T. equi* or *A. phagocytophilum*, which is consistent with the percentage determined at the scale of the whole Occitanie region where this tick species is established (11.8%) (Joly-Kukla et al. 2024). While recent spatial studies in the Occitanie region showed that some other pathogens, such as *Anaplasma marginale, Ehrlichia minasensis* and West Nile virus, were detected in *H. marginatum* (Joly-Kukla et al. 2024; Bernard et al. 2024a), these pathogens were not detected in our temporal study. All these data emphasize that combined spatio-temporal studies are needed to better understand the tick-borne pathogen ecology.

### 4.2 Seasonal dynamics of *Hyalomma marginatum*-borne pathogens

Ticks were recurrently infected with at least one pathogen through the collection period. Even though infection rates varied between the different pathogens, they were globally all detected recurrently from February to September. As already observed and mentioned for other tick species (i.e.(Lejal et al. 2019b)), our findings confirm that sporadic *H. marginatum* collections are not sufficient to assess the *H. marginatum*-borne pathogen prevalences reinforcing the importance of regular monitoring and underlying the necessity to keep preventing populations about the tick risk. In addition, the infectin rates were variable through the year with fewer infected ticks in spring compared to summer and winter. This finding might incite to be more cautious concerning the tick risk only at specific periods of the year, especially given that the tick densities are not similar across seasons. However, the fact that our findings have been obtained from a one-year study calls for caution in attempts to draw conclusions about this observed pattern and a multi-year survey would be necessary to assess a potential inter-annual recurrence of this pattern. In the literature, a three-year survey of *Ixodes ricinus-* borne pathogens showed fluctuations of the pathogen dynamics while similar patterns were observed for abiotic conditions from one year to another. In this case, the pathogen dynamics may be linked to the host dynamics (Lejal et al. 2019a) rather than abiotic conditions. Concerning *R. aeschlimannii* in *H. marginatum*, we can exclude the influence of the tick host on its dynamics due to maternal transmission and the expected status of the bacterium as a secondary symbiont (i.e.(Joly-Kukla et al. 2024)). A several-year survey of *R. aeschlimannii* in a sentinel site would therefore allow to focus on the exploration of the role of abiotic conditions on the dynamics of this bacteria.

In addition to the monthly survey in our current sentinel site of Pompignan, it would also be interesting to investigate another site, especially located in another geographic area. Indeed, it was reported that *R. aeschlimannii* infection rates significantly differ between sites from one geographic cluster to another in the Occitanie region, with higher infection rates for example in the Aude/Pyrénées-Orientales area (92.3% in average) compared to those estimated from the Hérault/Gard area (78.6% in average) (**Figure 1**) (Joly-Kukla et al. 2024). In this study, while fluctuations of infection rates were observed for all the detected pathogens, the most marked temporal variations were determined for *R. aeschlimannii*. A monthly survey in a sentinel site located in the Aude/Pyrénées-Orientales area would definitely be interesting to characterize the *R. aeschlimannii* seasonal dynamics in this second site and assess the potential recurrence of the temporal pattern observed in Pompignan (Hérault).

The presence of *R. aeschlimannii* in ticks varied significantly with the season. These temporal trends followed the same pattern regardless of the employed detection technique and targeted gene were very similar, thus reinforcing the robustness of this result. While its presence varied with the season, it was not interestingly the case for its loads. These contrasted results might probably be explained by the high variability in *R. aeschlimannii* loads observed between ticks collected during the same month and emphasize the importance of increasing the sampling effort as far as possible. However, as previously reported (Joly-Kukla et al. 2024), bacterial loads were higher in females than males. This could be due to the absence of ovaries in males, where *R. aeschlimannii* might predominantly be located since it is maternally transmitted (Joly-Kukla et al. 2024; Matsumoto et al. 2004; Azagi et al. 2017). This hypothesis might also explain the higher global infection rates in females compared to males. Additional information regarding the *R. aeschlimannii* location within the different tick organs would enable the formulation of a more precise hypothesis about its potential function in *H. marginatum*. Subsequently, its presence in the salivary glands could be determined, which would raise questions about its transmission to the tick host.

The status of *R. aeschlimannii* detected in *H. marginatum* was questioned in a recent study that demonstrated its high infection rate (87.3% in average) and its contrasted spatial distribution in the Occitanie region (Joly-Kukla et al. 2024). A maternal transmission coupled with high infection rates represent key characteristics to hypothesize that *R. aeschlimannii* might be a secondary symbiont in *H. marginatum* (Joly-Kukla et al. 2024; Bernard et al. 2024a). Based on the definition of secondary symbionts (i.e. (Su, Zhou, and Zhang 2013)), such an hypothesis suppose bacterial functions not necessarily required for the host survival but important for example for the defence against natural enemies (Oliver et al. 2010; Su et al. 2014) or to mediate the thermal tolerance of their host (Dunbar et al. 2007). Considering hypothetically that *R. aeschlimannii* is effectively a secondary endosymbiont for *H. marginatum*, its functional role for the tick remains currently unknown. In our study, even though transcriptomic and genomic data are missing to get more accurate information about the real status (pathogen vs endosymbiont) and the functional role of *R. aeschlimannii* in *H. marginatum*, the fact that estimated infection rates were very variable over time in both males and females with the same temporal patterns firstly reinforces the hypothesis of a secondary symbiont. Indeed, as observed in our results, a primary endosymbiont such as *Francisella*-like endosymbiont in *H. marginatum* would have been highly present in almost all the collected ticks (279 of the 281 analysed ticks) and its infection rates would not vary with time due to its obligatory characteristics for the tick’s survival. In addition, we observed a high increase in *R. aeschlimannii* infection rates in all the analysed ticks from June to the end of summer compared to winter and spring. In parallel to this observation, we interestingly demonstrated a significant correlation between *R. aeschlimannii* infection rates and the temperature. Based on these complementary results, we hypothesize that *R. aeschlimannii* might be involved in the stress response after an increase in temperature in the tick environment and might thus mediate the thermal tolerance of *H. marginatum*. Nevertheless, future experimental investigations should be performed in order to test this hypothesis. It would be indeed interesting to estimate both the *R. aeschlimannii* infection rates and loads from ticks experimentally placed in cages located in a shadow-spot vs a sunny-spot in the field, with a continuous monitoring of temperature. In addition, a genome analysis of this bacterium isolated from our *H. marginatum* ticks or the analysis of the relative expression of some functional genes involved in stress responses to temperature might also help us to validate or refute this hypothesis.

*T. equi* infection rates varied from 2.9% in April to 25.0% in August, although no significant differences were observed at the season scale. Within the same season, infection rates were variable between months, for example in summer (6.5% in June, 9.1% in July and 25% in August). While *T. equi* and *R. aeschlimannii* were detected every month, *A. phagocytophilum* was only detected in March, and during summer between June and September. These results highlight one more time the importance of a regular temporal monitoring of pathogens insidiously circulating and which may not be detected by a unique sampling at a given time of the year. Concerning *A. phagocytophilum*, the descriptive analysis interestingly revealed low infection rates before summer (4.5% in average in winter and 0% in spring) and much higher infection rates in summer (7.8%). Horses are not known to be susceptible to this bacterium. By contrast, birds, one of the main hosts of *H. marginatum* immatures, are susceptible to *A. phagocytophilum* infection (Bernard et al. 2024a) and would be able to transmit this bacterium to ticks during their two first life stages. The temporal variability observed for *A. phagocytophilum* might consequently result from the dynamics of immature stages and their associated hosts that maintain this bacterium at low prevalence. Because *A. phagocytophilum* is the causative agent of granulocytic anaplasmosis in horses, dogs, and humans, our findings demonstrate that it is important to be aware of specific periods during which there is a higher susceptibility to be bitten, in July for example (15.2% of analysed ticks were positive).

## 5 Conclusion

This study characterized the *H. marginatum*-borne pathogen composition and dynamics over eight months in 2022 in a sentinel site. These data are crucial for identifying monthly/seasonal patterns which could improve our understanding of *H. marginatum*-borne pathogen ecology and prevent *H. marginatum*-borne diseases. Our findings allowed to (i) identify pathogens already known to circulate in the study area; (ii) highlight strong temporal patterns at the month and season scales especially for *R. aeschlimannii*; and finally (iii) provide clues for venturing hypotheses on the potential function of *R. aeschlimannii* in *H. marginatum*, that could be involved in stress response due to the increase in temperature and potentially mediate the thermal tolerance of *H. marginatum*.

## Ethical approval

Not applicable.

## CRediT authorship contribution statement

**Conceptualization:** TP, SM, CJK. **Data curation**: CJK, TP. **Formal analysis**: CJK. **Funding acquisition**: TP. **Investigation**: TP, SM. **Methodology**: CJK, CG, DB, CR, SM. **Project administration**: TP. **Resources**: CJK, FS, MD, CG, SM, TP. **Software**: CJK, **Supervision**: TP, SM. **Validation**: CJK, FS, MD, CG, SM, TP. **Visualization:** CJK. **Roles/Writing - original draft**: CJK. **Writing - review & editing**: CJK, SM, TP.

## Acknowledgments

The authors are grateful to Maud Marsot and Facundo Munoz for their helpful advice on statistics.

## Supplementary data

**Figure S1:**
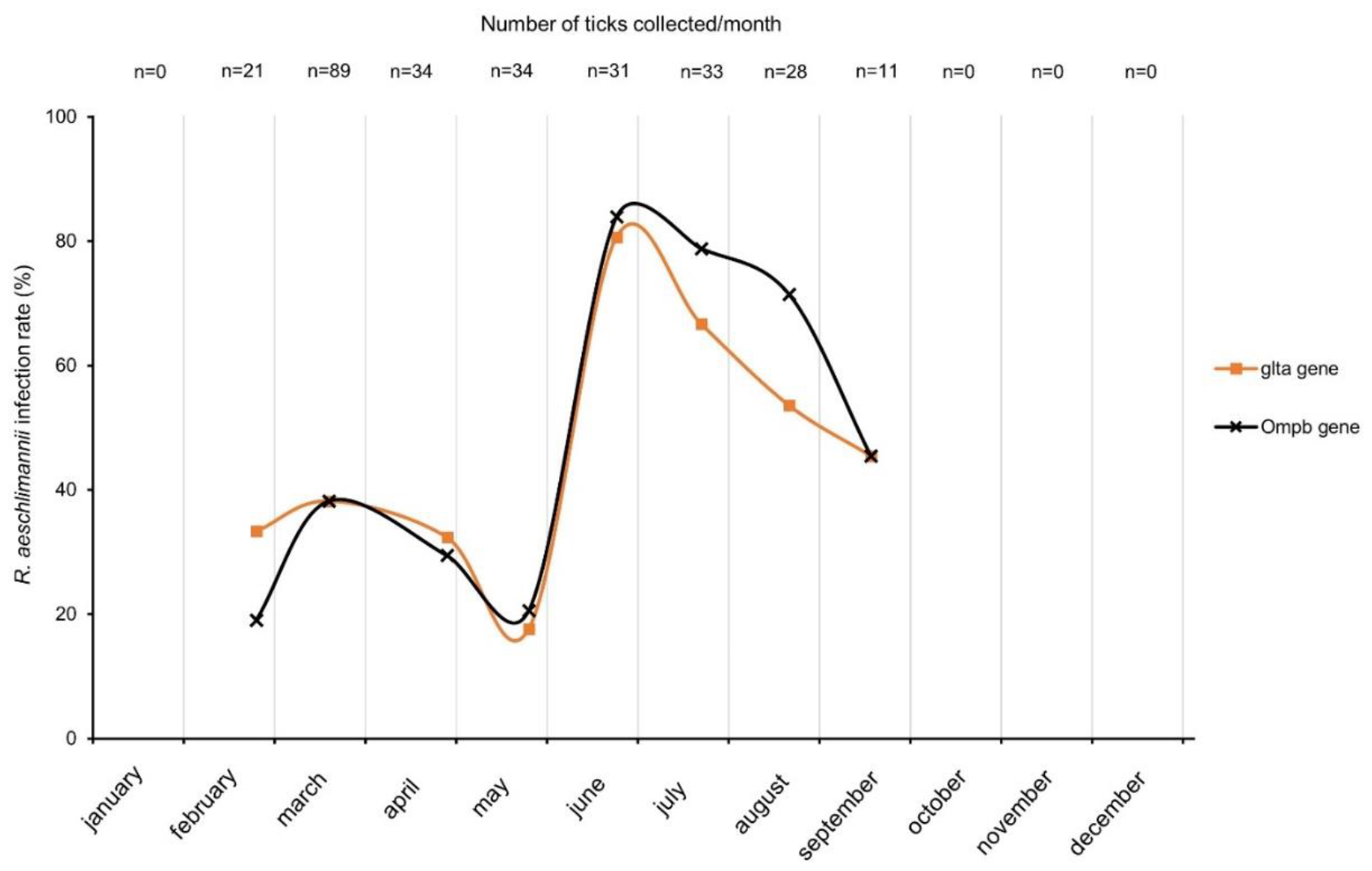
Monthly trends of *R. aeschlimannii* infection rates (%) obtained by either the BioMark™ assay targeting the *glta* gene (orange curve) or by qPCR that targets the *OmpB* gene (black curve). Dots correspond to the dates of collection. Winter: February to March; Spring: April to May; Summer: June to September). The number of collected ticks (abbreviation n) per month is indicated above the figure and below the figure for the seasons.

**Figure S2:**
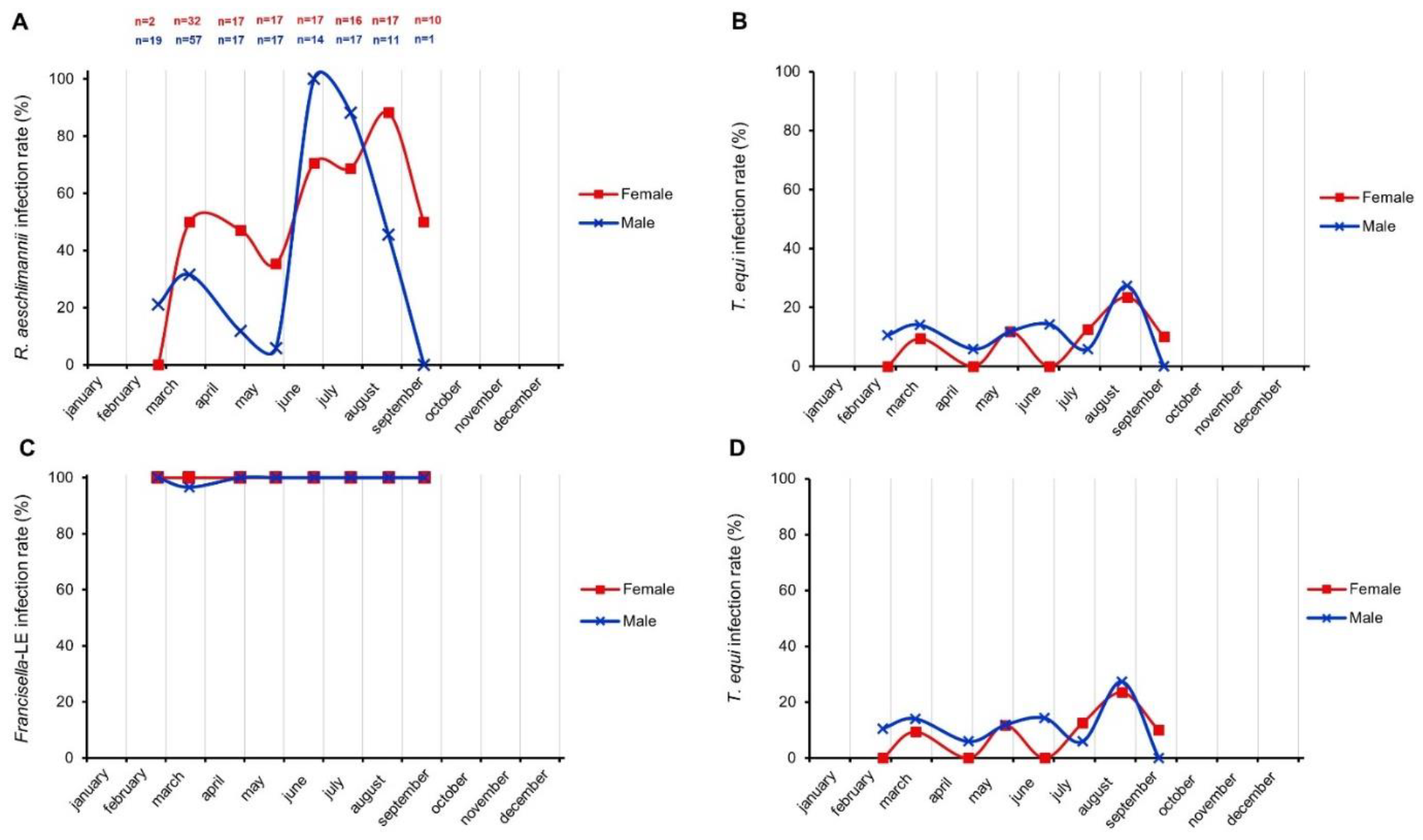
Monthly trends of infection rates (%) of **A**. *R. aeschlimannii*; **B**. *T. equi*; **C**. *Francisella*-LE and **D**. *A. phagocytophilum*, according to the tick sex. Dots correspond to the dates of collection. Winter: February to March; Spring: April to May; Summer: June to September. The number of female (red) and male (green) ticks analysed for each month is indicate above **A**. and is the same for **B**., **C**. and **D**.

## Data availability

Nucleotidic data were deposited in GenBank under the accession number PP663276. Scripts and code used for bioinformatics and statistical analysis are available on xxx and related project website.

